# Crowding induced morphological changes in synthetic lipid vesicles determined using smFRET

**DOI:** 10.1101/2022.05.31.494132

**Authors:** Steven D. Quinn, Lara Dresser, Sarah Graham, Donato Conteduca, Jack Shepherd, Mark C. Leake

**Affiliations:** School of Physics, Engineering and Technology, University of York, York, UK. YO10 5DD; York Biomedical Research Institute, University of York, York, UK. YO10 5DD; Department of Biology, University of York, York, UK. YO10 5DD

**Keywords:** Single-molecule, TIRF, FRET, membrane mechanics, lipid vesicle, molecular crowding

## Abstract

Lipid vesicles are valuable mesoscale molecular confinement vessels for studying membrane mechanics and lipid-protein interactions, and they have found utility among bio-inspired technologies including drug delivery vehicles. While vesicle morphology can be modified by changing the lipid composition and introducing fusion or pore-forming proteins and detergents, the influence of extramembrane crowding on vesicle morphology has remained under explored owing to a lack of experimental tools capable of capturing morphological changes on the nanoscale. Here, we use biocompatible polymers to simulate molecular crowding *in vitro*, and through combinations of FRET spectroscopy, lifetime analysis, dynamic light scattering and single-vesicle imaging, we characterize how crowding regulates vesicle morphology. We show that both freely-diffusing and surface-tethered vesicles fluorescently tagged with the DiI and DiD FRET pair undergo compaction in response to modest concentrations of sorbitol, polyethylene glycol and Ficoll. A striking observation is that sorbitol results in irreversible compaction, whereas the influence of high molecular weight PEG-based crowders was found to be reversible. Regulation of molecular crowding allows for precise control of vesicle architecture *in vitro*, with vast implications for drug delivery and vesicle trafficking systems. Furthermore, our observations of vesicle compaction may also serve to act as a mechanosensitive readout of extramembrane crowding.

## Introduction

Native biological membranes are highly complex and heterogeneous in both size and composition, motivating the development of controllable model-membrane systems whose physical and chemical parameters can be easily tuned^1^. A particularly valuable class of such systems are spherical synthetic vesicles, which comprise a phospholipid bilayer surface whose radius of curvature can be carefully controlled. Small unilamellar vesicles (SUVs), for example, range from *ca*. 10-100 nm in size, whereas large unilamellar vesicles (LUVs) and giant unilamellar vesicles (GUVs) are *ca*. 100 - 1000 nm and > 1 µm in diameter, respectively^1, 2^. Importantly, their phospholipid compositions can be tailored to enable variations in charge, localized membrane roughness, phase separation behaviour across the bilayer and membrane fluidity^1^.

Synthetic vesicles have enabled new insights into membrane mechanics^3^, interactions between lipids and proteins^4,5^ fusion dynamics^6^, and inspired the development of drug delivery vehicles due to their low toxicity, high loading capacity and controllable release kinetics^2-3^. LUVs have found particular utility as nanoscale containers for constraining biomolecules for single-molecule analysis, where inducing porosity into their membrane facilitates buffer exchange without removing the biomolecule under interrogation^4^. Additionally, vesicles can be biochemically programmed to interact, enabling lipid mixing and content exchange^5-7^, and the use of perturbative detergents, which alter their morphology^8-9^, are important in the context of lysis and for triggering release of encapsulated molecules^10^. It is also now clear that the structure and dynamics of synthetic vesicles are influenced by factors such as molecular crowding but new experimental approaches are required to explore these interactions further.

The living cell’s interior is a densely crowded environment, with up to 40 % of the cytoplasm occupied by solubilised macromolecules^11^. In this tightly filled space, excluded volumes give rise to steric repulsions, depletion attractions, reduced translational degrees of freedom, biomolecular shape changes, and diffusional effects, all of which contribute to the cell’s overall function^12^. In this context, the effect of macromolecular crowding on protein^13-14^ and nucleic acid^15-17^ structures have been heavily studied *in vitro*. For example, molecular crowding influences protein stability^18-19^, interactions, kinetics, diffusion^20^ and liquid-liquid phase separation events^21^. These results not only lend support to the idea that molecular crowding is a regulator of biomolecular activity, but that living cells may actively regulate crowding to enhance or adjust key processes^22^. Several different crowding mechanisms have been proposed depending on the biomolecule and co-solute, but recent theoretical studies and energy transfer experiments point towards an intriguing size dependence, where smaller molecular weight crowders are more effective than larger polymers^17, 23-24^. Despite such developments however, the question of how macromolecular crowding influences the global structure of single lipid vesicles remains largely under explored.

Initially, experiments assessing the impact of molecular crowding on the membrane involved the use of vesicles *in vitro* under relatively dilute concentrations of ions or other chemical factors. For example, crowding was observed to give rise to concentration- and polymer-dependent osmotic pressures and non-specific depletions resulting from effects assigned to excluded volumes^25-26^. When high molecular weight crowding agents were introduced, the preferential exclusion of macromolecules from the membrane bilayer introduced an osmotic imbalance which in turn altered the global membrane conformation and promoted membrane fusion^27-28^. Recent studies have shown that encapsulation of crowding agents within single vesicles leads to depletion forces that results in variations in membrane topology and changes to the surface area^29,30^. When highly hygroscopic crowders such as polyethylene glycol (PEG) were encapsulated, the membrane dehydrated, effectively leading to vesicle compression^31^. Optical microscopy experiments on 20-60 μm sized GUVs have also demonstrated that the encapsulation of high molecular weight PEGs induce membrane stress, oscillations in vesicle size, changes to the membrane tension, permeabilization and variations in the spatial orientation of membrane-bound molecules^32^. Furthermore, recent investigations on GUV crowding by nucleic acids point towards the formation of local elastic deformations and transient instabilities in the membrane^33^. These results support a model in which molecular crowding influences the architecture, dynamics and integrity of biological membranes. However, optical microscopy experiments on GUVs, such as those reported, only reveal macroscopic changes taking place within a 2-dimensional image plane, without quantitatively reporting on molecular level changes across the entire volume. Moreover, it is currently unclear what the influence of low molecular weight crowders are. These challenges, combined with the need to assess the structural integrity of vesicles at the opposite end of the membrane-curvature space, motivated us to extend our single-molecule Förster resonance energy transfer (smFRET) toolbox to quantitatively assess conformational changes taking place within sub-micron sized LUVs in response to a range of molecular crowders *in vitro*.

Here, we employed a single-vesicle assay based on measuring the extent of smFRET between lipophilic fluorophores integrated into the membrane of LUVs to evaluate their conformation in response to crowding agents in the extravesicle space. We first integrated the probes into the LUV bilayer and observed conformational changes in response to crowding via ensemble fluorescence spectroscopy and time-correlated single photon counting. Electron microscopy and dynamic light scattering approaches were then used to quantify structural variations, before wide-field total internal reflection fluorescence microscopy was used to capture the FRET response from single vesicles. By monitoring changes to the FRET efficiency within freely-diffusing and surface-immobilized vesicles, we correlate crowding-induced changes in fluorescence signals to morphological changes within single vesicles. In particular, we found that both sorbitol, a model sugar based cosolvent for low molecular weight crowding^34^, and high molecular weight crowders such as PEG400, Ficoll400 and PEG8000 could induce enhancements to the observable FRET efficiency. Compaction induced by the high molecular weight crowders was found to be reversible, whereas sorbitol-induced vesicle compaction was permanent. We expect the presented tools will be widely applicable beyond the interactions studied here and we discuss the implications of our findings for enabling control over vesicle morphology, for drug delivery, vesicle trafficking and the regulation of vesicle curvature.

## Materials and Methods

### Lipid vesicle preparation

1-palmitoyl-2-oleoyl-glycero-3-phosphocholine (POPC) and 1-palmitoyl-2-oleoyl-sn-glycero-3-phospho-L-serine (POPS) lipids in chloroform were purchased from Avanti Polar Lipids and used without any additional purification. 1,1’-Dioctadecyl-3,3,3’,3’ Tetramethylindocarbocyanine Perchlorate (DiI) and 1,1-Dioctadecyl-3,3,3,3-tetramethylindodicarbocyanine (DiD) were obtained from ThermoFisher Scientific. Synthetic vesicles were prepared via the extrusion method as previously described^8-9^. Briefly, mixtures of lipids and membrane stains were mixed in chloroform at final lipid concentrations of 10 mg lipid/ml. The solvent was then evaporated by nitrogen flow to create a dry lipid film, subsequently hydrated in 50 mM Tris buffer (pH 8.0) and mixed by vortex. The resuspended solution was then extruded through a polycarbonate membrane filter to produce vesicles of appropriate diameter.

### Ensemble Fluorescence Spectroscopy

Fluorescence emission spectra obtained from DiI and DiD loaded vesicles in 50 mM Tris buffer (pH 8) were recorded using a HORIBA Fluoromax-4 spectrophotometer with λ_ex_ = 532 nm. All experiments were performed using a final lipid concentration of 25 μM. Apparent FRET efficiencies, *E*_*FRET*_, were estimated via *E*_*FRET*_ *= (I*_*A*_*/[I*_*A*_*+I*_*D*_]), where *I*_*A*_ and *I*_*D*_ are the integrated, background-corrected fluorescence emission intensities of the donor, DiI, and acceptor, DiD, respectively. Data points plotted represent the mean and standard deviation obtained from three separated experimental runs.

### Time Correlated Single Photon Counting

Time-resolved fluorescence decays were collected using a FluoTime300 time-correlated single photon counting spectrophotometer equipped with a hybrid PMT detector (Picoquant, Germany). Decays were measured under magic angle conditions using pulsed excitation of 532 nm at 80 MHz, and emission of 565 nm for DiI-DiD loaded vesicles. Excitation of 485 nm (20 MHz) and emission of 600 nm was used for vesicles incorporating the tension reporter FliptR^35^. All experiments were performed using a final lipid concentration of 25 μM in 50 mM Tris buffer (pH 8). Decays were acquired until 10^4^ counts at the decay maximum were observed and fitted by iterative re-convolution of the instrument response function and the observed fluorescence decay using a multi-exponential decay function of the form 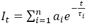, where *I*_*t*_ is the intensity at time, *t*, and *t*_*i*_ and *a*_*i*_ represent the fluorescence lifetime and fractional amplitude of the *i*^th^ decay component.

### Dynamic Light Scattering (DLS)

The hydrodynamic diameter, *d*_*H*_, of freely diffusing vesicles was estimated using a Zetasizer μV system equipped with a *λ*_o_ = 633 nm wavelength line. Briefly, Brownian motion of LUVs in solution gives rise to fluctuations in the intensity of back scattered light at *θ* = 178°. This was used to produce a correlation function, *G(τ)*, via *G*^*2*^*(τ) = <I(t)I(t+τ)>/<I(t)*^*2*^*>* where *τ* is the lag time. Correlation curves were fitted to a model of the form 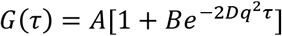 where *D* is the vesicle diffusion coefficient, *A* and *B* are positive constants and 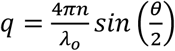, where *n* is the refractive index of the solution (*n*=1.33). Hydrodynamic diameters were then calculated according to the Stokes-Einstein relationship^36^ and reported as the mean and standard deviation from three separated experimental runs.

### Scanning Electron Microscopy (SEM)

SEM micrographs of vesicles non-specifically bound to a silicon substrate were acquired using a JEOL JSM-7800F system operating at 5 kV ^5,37^. Vesicles were prepared in 50 mM Tris buffer (pH 8) containing molecular crowders at concentration specified in the main text, diluted ∼10x in deionized water and vortexed. The vesicles were then applied to the silicon substrate and the solution evaporated. The substrate was then sputtered with a 5 nm thick Pt/Pd layer to avoid charging effects and possible damage of the vesicles during the micrographs acquisition. Vesicle diameters were then determined using ImageJ, and histograms produced using 25 nm bin widths.

### Cryo-Transmission Electron Microscopy (Cryo-TEM)

Cryo-TEM was used for direct visualization of vesicle bilayers. To prepare samples for cryo-TEM analysis, Quantifoil copper R 1.2/1.3 200 Mesh grids (Electron Microscopy Sciences) were prepared by glow discharging at 20 mA and 0.26 millibar for 1 minute in a Pelco easiGlow system. Small volumes (2 µL) of the vesicle sample was applied to the carbon side of the EM grid in 90% humidity, then liquid was blotted off for 0.5 s and the grids plunge-frozen into precooled liquid ethane using a Vitrobot system (Thermo Scientific). This process enabled single vesicles to be embedded within a thin layer of amorphous ice, preserving them in their native state. The samples were then evaluated using a Thermo Scientific Glacios Cryo-TEM electron microscope. TEM images were acquired using an accelerating voltage of 200 kV, and 120,000x magnification. Vesicle sizes were then measured using ImageJ.

### Total Internal Reflection Fluorescence (TIRF) microscopy

Microfluidic flow cells were constructed as described previously^38^ and sequentially incubated with 0.1 mg/mL BSA-Biotin, 1 mg/mL BSA and 0.2 mg/mL NeutrAvidin. After each incubation step the flow cells were rinsed with buffer (50 mM Tris, pH 8) to remove unbound material. Biotinylated vesicles containing 0.1 mol % DiI and 0.1 mol% DiD were then added to the surface using a final concentration of 70 μg/mL lipids in imaging buffer (50 mM Tris, 6 % (W/V) D-(+)-glucose containing 1 mM Trolox, 6.25 μM glucose oxidase and 0.2 μM catalase) and incubated for 15 minutes at room temperature to achieve a surface coverage of ∼150-200 vesicles per 50 × 50 μm field of view. After incubation, the flow cells were rinsed with imaging buffer. TIRF microscopy was then performed on a custom-modified inverted microscope (Nikon Eclipse Ti) containing a CFI Apo TIRF 100 x NA 1.49 oil-immersion objective lens (Nikon). TIRF illumination was provided by a TEM_00_ 532 nm wavelength line (Obis, Coherent) at < 8.2 mW cm^-2^. Emission was separated from the excitation line via a dichroic and emission filter mounted beneath the lens turret (Chroma 59907-ET-532/640). DiI and DiD emission was then spatially separated using a DualView image splitter (Photometrics) containing a dichroic filter (T640LPXR, Chroma) and band pass filters (ET585/65M and ET700/75M, Chroma) and imaged in parallel on a back-illuminated Prime 95B CMOS camera cooled to -30°C (Photometrics). After each addition of crowder in imaging buffer, 500 frame movies were acquired with 50 ms exposure time. Recorded images were then analysed in MATLAB (R2019a) using iSMS single-molecule FRET microscopy software^39^. Briefly, the donor and acceptor emission channels were aligned, and background-corrected DiI and DiD integrated emission trajectories were obtained within the excitation field. Apparent FRET efficiencies across the trajectories were then calculated using *I*_*A*_*/(I*_*A*_*+I*_*D*_) as previously described, and related to the mean distance between probes, 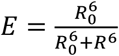, where *R*_*o*_ is the Förster radius (5.3 nm)^8^. *E*_*FRET*_ histograms from *N* > 2000 vesicles were then produced using bin widths of 0.01.

## Results and Discussion

### Sorbitol induces conformational changes in freely diffusing lipid vesicles

We first used smFRET between lipophilic membrane stains to explore the structure of LUVs composed of POPC lipids in the presence of synthetic molecular crowding agents. Previous studies evaluating membrane deformation have largely relied on encapsulating molecular crowders or introducing pegylated lipids into GUV bilayers, with single-colour fluorescence imaging used to evaluate macroscopic changes^40^. However, such approaches do not provide detailed information at the molecular level, nor do they report on vesicles smaller than the optical diffraction limit^8^. We therefore applied smFRET to quantify nanoscale structural variations in LUVs of ∼200 nm in size in response to dilute crowding conditions.

We first optimized the amount of donors (DiI) and acceptors (DiD) per LUV (1:1 ratio, 0.1 mol % of each dye), such that the average FRET efficiency (*E*_*FRET*_) per vesicle was initially ∼0.5. This corresponds to an average DiI-DiD separation close to their Förster radius, and allows for nanoscale changes because of vesicle compaction or swelling, to be quantified by an observable increase or decrease in E_FRET_. To mimic low molecular weight crowding, we used the polyol osmolyte sorbitol, an established low molecular weight crowding agent which has previously been applied to regulate protein clustering^41-43^, induce nuclear organization and compact chromatin^44^.

Dynamic light scattering first confirmed the formation of labelled vesicles with a log-normal hydrodynamic diameter centred on 227 nm (**Figure S1**). When vesicles were prepared without labels, a similar distribution was observed, providing confidence that the labelling process does not perturb morphology (**Figure S1**). We then recorded the ensemble fluorescence spectra obtained from the labelled vesicles in solution. As shown in **Figure 1A**, DiI and DiD loaded LUVs displayed a progressive increase in sensitized acceptor emission and E_FRET_ as the sorbitol concentration progressively increased, translating to a reduction in the mean DiI-DiD separation distance from 5.02 ± 0.02 nm in the absence of crowding to 4.76 ± 0.01 nm in the presence of 3M sorbitol (**Figure 1B**). To further confirm the presence of an energy transfer mechanism, we measured the amplitude weighted fluorescence lifetime, τ_av_, of DiI in the presence of DiD via time-correlated single photon counting. Here, *τ*_*av*_ decreased with increasing sorbitol, consistent with a progressive quenching of DiI and corresponding increase in E_FRET_ (**Figure 1C**). The decays fitted well to a bi-exponential decay after reconvolution with the instrumental response function, which we assigned to energy transfer between dyes on the inner and outer leaflets. Indeed, a bootstrap analysis revealed that both lifetime components, termed *τ*_*1*_ and *τ*_*2*_, progressively decreased with increasing sorbitol (**Figure S2**), suggestive of global conformational changes on both sides of the membrane. Taking these measurements together, the data pointed towards fluorophore packing in the ensemble and were suggestive of vesicle compaction.

**Figure 1.**
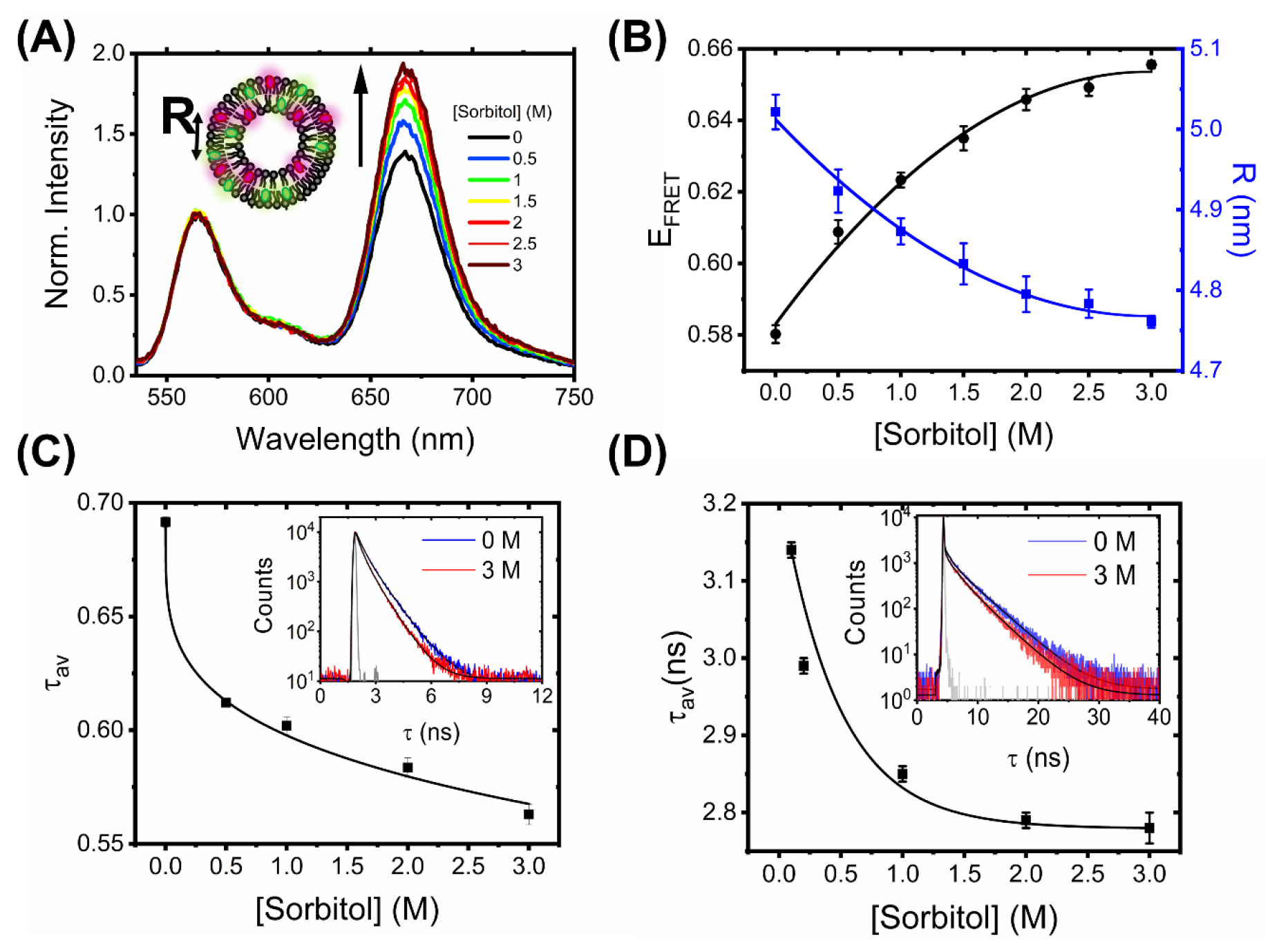
Sorbitol induces conformational changes in freely diffusing vesicles. (A) Normalized fluorescence emission spectra of DiI-DiD LUVs in the absence and presence of sorbitol (λ_ex_ = 532 nm). Inset: schematic illustration of the vesicles, where *R* corresponds to the mean dye-dye separation distance. (B) Corresponding variations in *E*_*FRET*_, and *R*. Heuristic fits shown are quadratic (solid black lines), determined with Python 3 using NumPy’s polyfit routine. (C) The amplitude weighted average lifetime of DiI and (D) FliptR as a function of sorbitol. Insets correspond to the time-resolved fluorescence decays in the absence (blue) and presence of 3M sorbitol (red). Solid black lines represent biexponential fits to the raw data and the solid gray lines represent the instrument response functions.

To test whether sorbitol induced variations in membrane tension, we next evaluated changes in the fluorescence lifetime of 1 mol % Fluorescent LIPid Tension Reporter (FliptR) incorporated within the LUV bilayer. The FliptR lifetime correlates well with membrane tension^35^ and in our case, we found that decays fitted well to a biexponential model. In line with previous observations, the longer lifetime component, *τ*_*2*_, only accounted for a small fraction of the overall signal. In the absence of sorbitol the ensemble FliptR lifetime was found to be 3.14 ± 0.01 ns, corresponding to a situation where the molecules reside in a planar conformation. However, as sorbitol was progressively titrated, we observed a reduction in lifetime to ∼90 % of its original value (**Figure 1D, Figure S3**), which we hypothesise is due to a fraction of FliptR molecules twisting into a conformation that is sterically more favourable because of changes to the lipid packing density and decreased membrane tension. Similar to observations made with single vesicles in response to surfactants^45^, we speculate that phospholipid bilayers mixed well with sorbitol leads to a situation where bilayer components are forced by entropy resulting in mixed sorbitol-lipid aggregates, local membrane undulations, and reduced membrane tension.

The ensemble FRET, lifetime and FliptR analysis broadly supports conformational rearrangements taking place in LUVs in response to sorbitol. However, to understand the parameters that affect this process, we next moved to interrogate the impact of vesicle size, composition and phase. When vesicles of various sizes (100, 400 and 1000 nm), as confirmed by DLS (**Figure S4**), interacted with sorbitol, the FRET efficiencies in all cases increased with crowder concentration, signifying similar dye-dye distance changes (**Figure S5**). Indeed, the relative magnitude of the FRET enhancement compared to those observed from 200 nm diameter vesicles was similar across all conditions tested.

To assess the impact of lipid composition and phase, we next probed the interaction between sorbitol and 100 nm diameter vesicles containing POPS lipids as a function of temperature and compared the relative change in FRET efficiency to similarly sized vesicles composed of POPC (**Figures S5**). POPC has a gel-to-liquid phase transition temperature, *T*_*M*_, of -2°C, and is therefore in the liquid phase at temperatures > 4°C, whereas *T*_*m*_ = 14°C for POPS, below which the vesicles are in the gel phase. We observed that the initial FRET efficiency magnitude was similar for both sets of vesicles at 4°C, 21°C and 37°C, however the relative change in E_FRET_ for each sorbitol condition was generally larger in the case of POPC vesicles. One possible explanation for the observed difference could rest in the hydration of the lipid carbonyls. POPS lipids are less mobile and give rise to dehydrated vesicle forms^46^, suggesting uptake of sorbitol into the bilayer may be key for conferring the observed changes.

### Morphological Characterization of Single Vesicles in Response to Sorbitol

To explore whether the observed FRET changes were coupled with changes in vesicle morphology, we next investigated the vesicle sizes using scanning electron microscopy (SEM). As shown in **Figure 2A**, vesicles composed of POPC lipids in the absence of sorbitol were mostly spherical (circularity = 0.87 ± 0.16), in line with previous observations^37^, with a size distribution centred on 132 ± 3 nm (FWHM = 125 ± 8 nm). However, with the addition of sorbitol, the peak shifted to 105 ± 5 (FWHM = 155 ± 15 nm) (**Figure 2B**) and the circularity reduced to 0.55 ± 0.12, consistent with a model in which sorbitol leads to local undulations in membrane morphology and an overall compaction. This analysis was further supported by variations in the vesicle hydrodynamic diameter (*d*_*H*_) reported by DLS, where the size distributions in solution progressively decreased upon sorbitol addition **(Figure S6)**. We note that the vesicle sizes reported by DLS are generally larger than those reported by SEM, likely due to vesicles being dehydrated and fixed under vacuum for SEM imaging. Nevertheless, both measurements pointed towards sorbitol-induced structural changes taking place within single vesicles.

**Figure 2.**
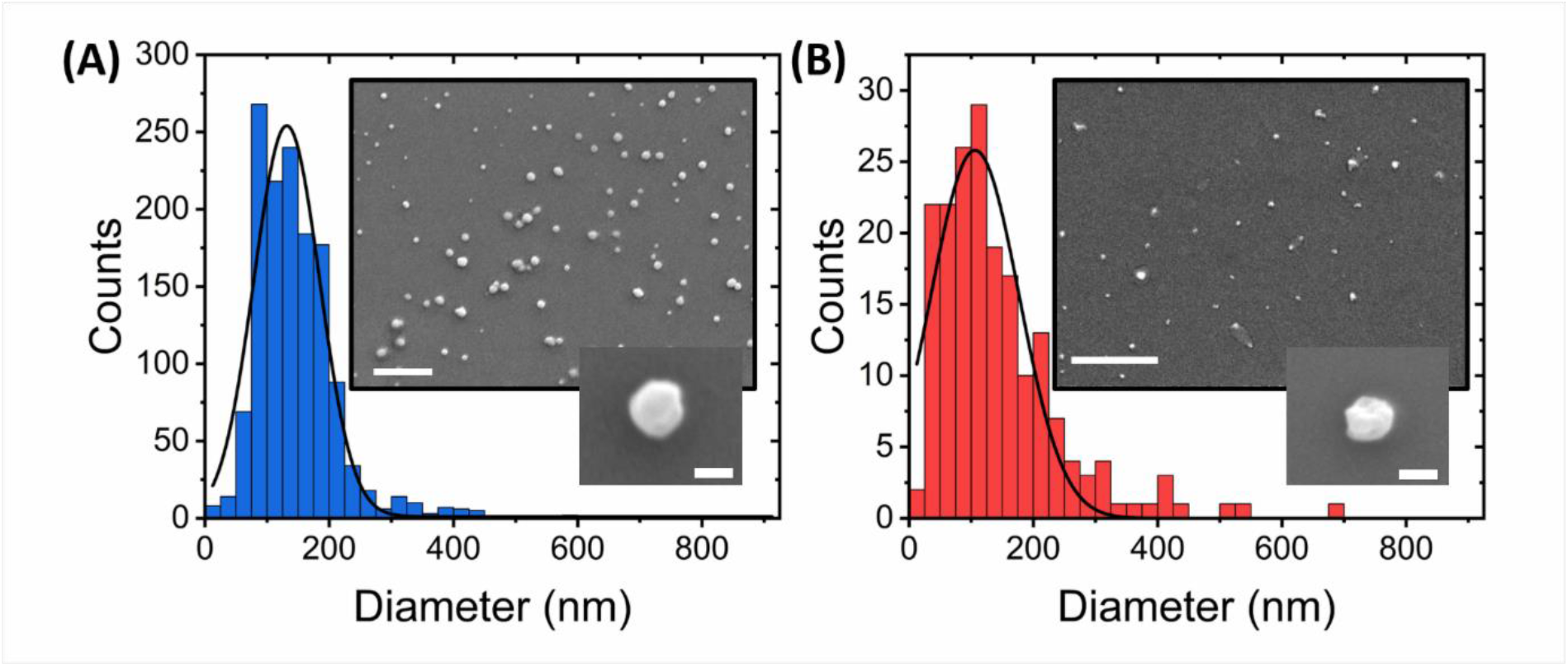
Sorbitol induces compaction and undulations in single lipid vesicles. Quantitative comparison of diameter distributions of POPC vesicles in (A) the absence and (B) presence of 3 M sorbitol. Insets: representative SEM images of immobilized vesicles under the respective conditions. Scale bars = 1 μm and 100 nm in the larger and smaller insets, respectively.

To further characterize the morphology of the vesicles, we also performed cryo-transmission electron microscopy (cryo-TEM). This enabled us to visualize the internal structures of hydrated vesicles in a frozen state, bypassing the requirement for dehydration. In the absence of crowding agents, the majority of vesicles were classified as unilamellar, exhibiting only a single bilayer **(Figure 3A-D)**, with lower fractions (< 20 %) containing double **(Figure 3E,F)** or multiple layers **(Figure 3G,H)**. All of the vesicles were intact, lacked any evidence of pore formation and in line with our SEM analysis, they were spherical in nature with diameters in the range 60-315 nm. Among the observed variants, we also observed larger vesicles containing encapsulated smaller vesicles **(Figure 3I)** unusually shaped structures in the form of a bowling pin **(Figure 3J, K)** and large conglomerates **(Figure 3L)**. Interestingly, the structures produced from our synthetic vesicle preparation have a striking resemblance to those observed from extracellular vesicles isolated from cerebrospinal fluid^47^, further validating our systems as excellent membrane mimetics. Of those vesicles classified as unilamellar, we observed a mean particle size of 157 ± 5 nm, with a membrane thickness of 6 ± 0.7 nm (*N* = 84) **(Figure S7**). In the presence of 0.5 M sorbitol, the observed species were structurally similar (**Figure 3 M, N, O)** with some vesicles displaying evidence of irregular membrane undulations and bulging **(Figure 3P)**. However, in line with our SEM and smFRET analysis, the vesicle size distribution shifted to smaller values and the unilamellar vesicle size reduced by ∼30 % to 115 ± 5 nm (*N* = 138). When 1 M sorbitol was introduced, most vesicles were intact (**Figure 3Q,R)**, however, others displayed signs of damaged, porous membranes, as indicated by the lack of a visible bilayer and lack of electron-dense material spaced across the projection of the vesicle surface **(Figure 3S)**, and the particle size reduced further to 82 ± 5 nm (*N* = 24). Based on this evidence, we suggest that the synthetic lipid vesicles studied in this work undergo global morphological changes upon sorbitol addition that broadly involves vesicle compaction coupled with membrane undulations, direct membrane damage, and the formation of pores.

**Figure 3.**
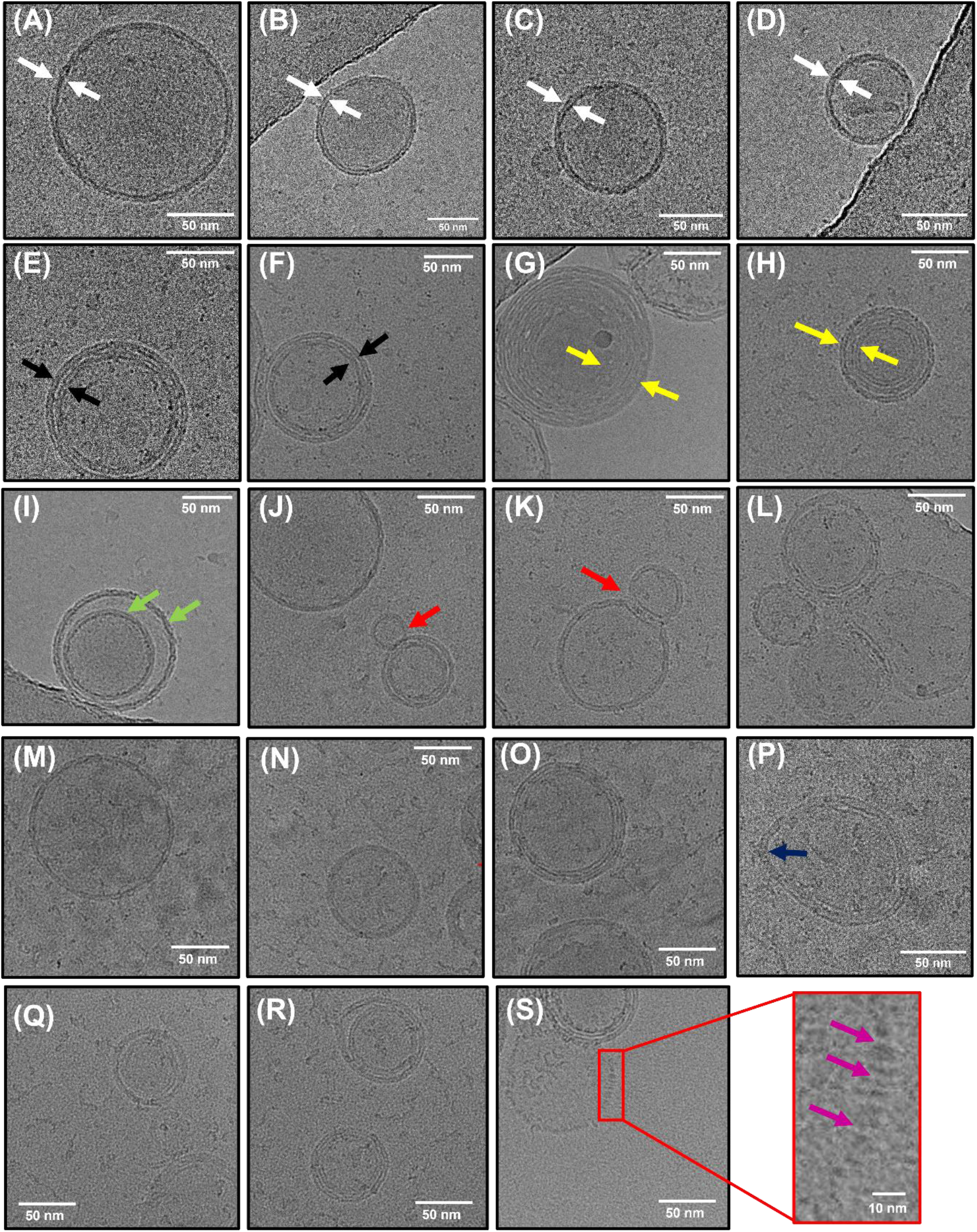
Morphological characterization of lipid vesicles by Cryo-TEM. Representative examples of (A-D) unilamellar (white arrows), (E-F) bilamellar (black arrows) and (G-H) multilamellar (yellow arrows) vesicles in the absence of sorbitol. Also shown are examples of (I) encapsulated vesicles (green arrows), (J-K) bowling pin shaped vesicles and (L) large conglomerates. (M-O) Representative examples of vesicles in the presence of 0.5 M sorbitol, and (P) example of vesicle displaying irregular bulging. (Q-R) Representative examples of vesicles in the presence of 1 M sorbitol with (S) example of broken membrane indicative of regions of broken membrane (purple arrows).

### Sorbitol-induced compaction of surface-tethered vesicles is irreversible

To further visualize the observed compaction events, we next evaluated the mean FRET response from single surface-tethered vesicles under crowding conditions via a custom-modified wide-field objective-type total internal reflection fluorescence microscope^47^. Prior to extrusion, the lipid suspension contained 1 mol % of biotinylated lipids (Biotin-PE), allowing formed vesicles to be tethered to a glass substrate via NeutrAvidin^9^. The application of picomolar vesicle concentrations to a NeutrAvidin coated surface led to the detection of 195 ± 14 FRET-active vesicles per 25 × 50 μm, with a mean nearest-neighbour vesicle separation distance of ∼1 μm (**Figure 4A**). In the absence of sorbitol, the vesicles displayed fluorescence across donor and acceptor emission channels, indicative of substantial FRET due to their close proximity. DiI and DiD intensity distributions obtained from *N* = 1,566 foci displayed lognormal behaviour (**Figure 4B**), which likely represents a distribution of vesicle sizes on the surface, consistent with our EM analyses^48,49^. As sorbitol was progressively added, we then recorded variations in the FRET efficiency per vesicle via changes to the DiI and DiD emission. Specifically, we observed a progressive increase in sensitized acceptor emission as crowding was increased because of enhanced FRET between the dyes (**Figure 4C**). During the titration, the number of foci per field of view and the mean total fluorescence intensity per vesicle defined as the sum of DiI and DiD emission intensities remained largely unchanged, providing confidence that sorbitol addition left vesicles intact on the surface. In the absence of sorbitol, the FRET efficiency distribution displayed Gaussian behaviour and was centred on 0.45, corresponding to a mean DiI-DiD separation distance of 5.4 nm. With increasing levels of sorbitol, the peak of the distribution then shifted by 15 % to 0.53 at 2.5 M, corresponding to a reduction in the mean DiI-DiD separation (**Figure 4D**). Given the total intensity of the foci remained largely invariant upon sorbitol addition, the positive shift in the FRET efficiency population distributions observed as sorbitol was added (**Figure 4E**) could not therefore be attributed to lipid loss or photophysical artefacts, but rather to structural alterations within single vesicles where the mean donor-acceptor separation distance, <*d*> progressively reduced. The crowding-induced morphological changes, observed here are thus consistent with compaction and the data is broadly complementary to previous observations where encapsulated molecular crowders led to vesicle bulging^32^. In our case the progressive addition of sorbitol led to instantaneous morphological changes on our measurement timescale. Furthermore, we observed that the compaction by sorbitol was irreversible. As shown in the top panel of **Figure 4F**, the vesicles were initially prepared in the absence of sorbitol and had a FRET distribution centred on 0.42. After incubation with 2.5 M sorbitol, a positive shift in the FRET distribution was observed (**Figure 4F, middle panel)**, however, after rinsing the flow cell with buffer, the distribution post-sorbitol addition remained unchanged (**Figure 4F, bottom panel**). This observation rules out an excluded volume effect as the underlying cause of the compaction since freely-diffusing sorbitol was removed from solution. Instead, this observation points towards a situation where after interaction of the membrane with sorbitol, the phase transition temperature is irreversibly depressed in a dose-dependent manner via an interaction that we speculate could involve sorbitol interactions with the functional phosphate groups on the lipids.

**Figure 4.**
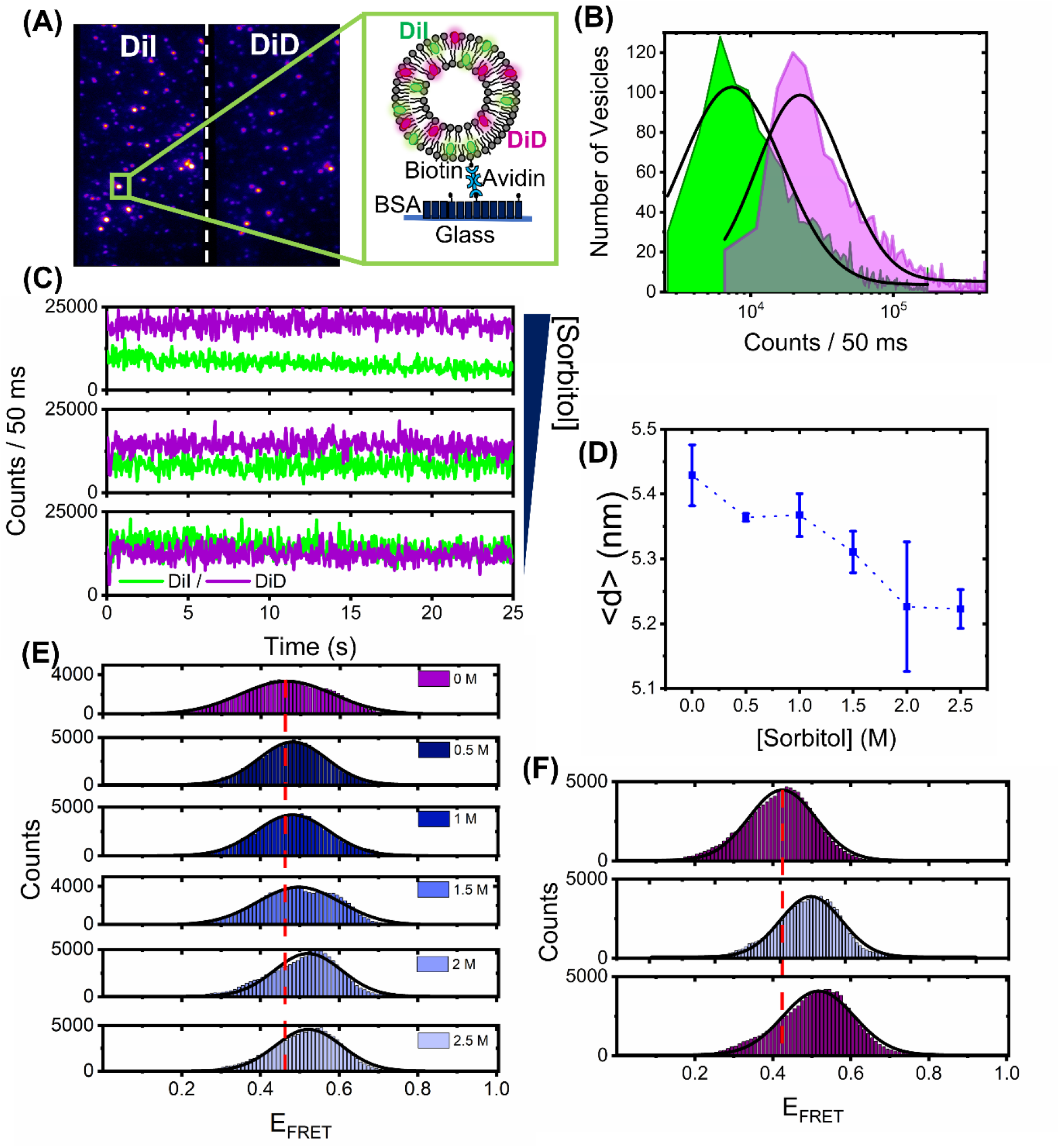
Sorbitol induces irreversible structural changes in single surface-tethered vesicles. (A) Representative wide-field TIRF image of surface tethered vesicles composed of DiI and DiD. Donor and acceptor emission channels are shown on the left- and right-hand side of the dashed line, respectively. Inset: surface immobilization scheme. Single vesicles containing biotinylated lipids are immobilized onto a BSA-Biotin coated glass coverslip via NeutrAvidin. (B) Fluorescence intensity population histograms of DiI (green) and DiD (magenta) obtained from surface tethered vesicles in the absence of sorbitol. (C) Representative time traces of DiI (green) and DiD (magenta) obtained from single surface tethered vesicles with 0 mM, 1 M and 3 M sorbitol. (D) Representative variation in the peak probe separation distance, <*d*>, as a function of sorbitol concentration. Also shown are fitting errors associated with the application of single Gaussian distributions to histograms of the probe-separation distance. (E) Corresponding variations in the FRET efficiency histograms obtained for *N* > 2,000 vesicles. (F) FRET efficiency histograms obtained from *N* > 2,000 vesicles at 0 mM sorbitol, after a 3 M sorbitol rinse step (middle panel), and after vigorous washing of the sample with imaging buffer (lower panel). The dashed red lines in (E) and (F) correspond to the peak positions of the FRET efficiency histograms obtained in the absence of sorbitol. Solid black lines represent single Gaussian fits.

Having established the DiI-DiD FRET pair as a sensitive indicator of vesicle compaction, we next explored the effect of higher molecular weight crowding agents known to induce excluded volume effects on biomolecular morphology. Here, we evaluated the fluorescence response of surface-immobilized vesicles labelled with DiI and DiD in response to variations of PEG and Ficoll. As reported previously, macromolecular crowding by PEG and Ficoll, even at modest concentrations in solution, can trigger substantial excluded volume effects, regulate mesoscale biological functions and impact the conformations of single biomolecules^15, 50-53^. We therefore hypothesised that high molecular weight crowders in the extravesicular space could influence the structural integrity of single lipid vesicles. By using the same immobilization strategy and conditions as described previously we imaged the vesicles under low-excitation TIRF conditions with 50 ms time integration before and after addition of molecular crowders. In the absence of crowding agents, the FRET efficiency distributions obtained from *N* > 2000 vesicles displayed Gaussian behaviour centred on 0.4 corresponding to <*d*> = 5.7 nm (**Figure 5 A-D**). When 5 – 20 % (*w/w*) of the low molecular weight crowder PEG 200 (a 200 Da grade of PEG) was injected, the distributions remained largely invariant, indicative of little-to-no effect on vesicle morphology (**Figure 5A**). However, when similar experiments were performed with the highly branched Ficoll 400 and linear PEG 400, both of which represent 400 Da crowding agents, we observed positive shifts in the population histograms, with <*d*> decreasing to 5.4 nm at 10-15 % (*w/w*) crowder (**Figure 5B, C**). When PEG 8000 was introduced, the FRET distribution shifted further, peaking 0.6 at 20 % (*w/w*), with <*d*> = 4.9 nm (**Figure 5D**). On the basis of spherical vesicle morphologies, the change in inter-dye distance observed from the vesicles in the presence of Ficoll 400, PEG 400 and PEG 8000 scales directly with the vesicle radius, and the change is thus supportive of compaction. The maximum FRET efficiency shift observed across the titration displayed a dependence on crowder size in the order PEG 8000 > PEG 400 > Ficoll 400 > PEG 200 **(Figure 5E)**, suggesting that high molecular weight polymers are more effective at inducing this morphological change. Although the exact nature of these variations requires further investigation, our data suggests that the DiI-DiD FRET pair is sensitive to the three-dimensional structure of the vesicle, and that changes to the FRET signal may arise from an excluded volume effect, with the higher molecular weight polymers leading to more pronounced changes in morphology. A particularly striking observation was that in the cases of PEG 400 and PEG 8000, the FRET efficiency distributions recovered to their original state after the vesicles were thoroughly washed with buffer solution (**Figure 5F**), indicating reversibility. Unlike in the case of sorbitol where we speculate that direct sorbitol-lipid interactions lead to membrane dehydration and irreversible compaction, here we speculate crowding arises primarily through an excluded volume effect. By removing the crowding agents from solution, we effectively remove this influence, and thereby allow the vesicles to recover to their original form, as schematically shown in **Figure 5G**.

**Figure 5.**
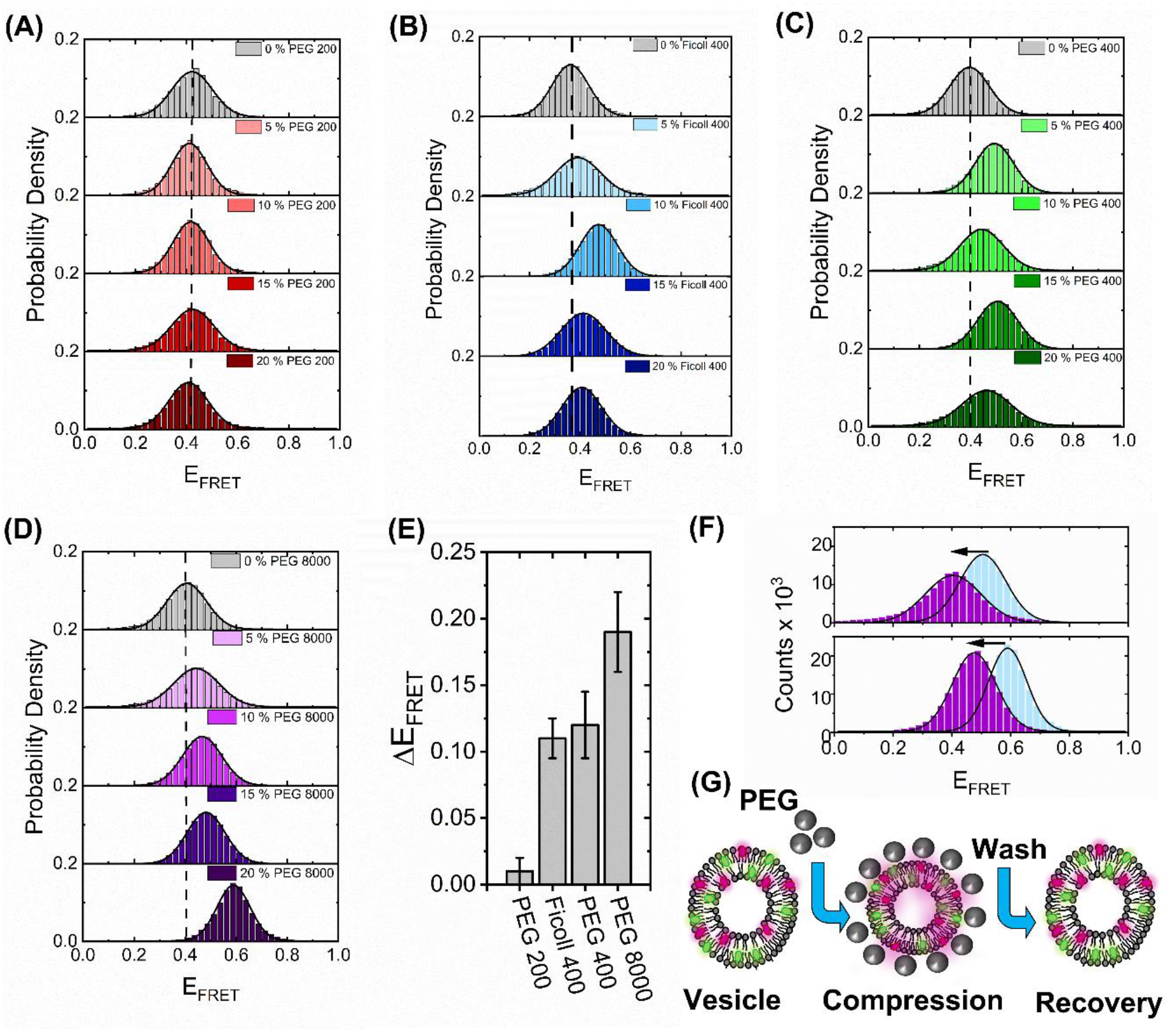
High molecular weight crowders induce reversible vesicle compaction. Representative variations in the FRET efficiency histograms obtained for *N* > 2,000 vesicles in the absence and presence of (A) PEG 200, (B) Ficol 400 and (C) PEG 400 and (D) PEG 8000 at 5 %, 10 %, 15 % and 20 % (*w/w*) in 50 mM Tris buffer (pH 8). Dashed lines correspond to the peak positions of the FRET efficiency histograms obtained in the absence of crowder. Solid black lines represent single Gaussian fits. (E) Comparative bar plot summarizing the maximum variation in FRET efficiency observed under different crowding conditions. (F) Representative FRET efficiency histograms obtained from *N* > 2,000 vesicles in the presence of PEG400 (blue, top panel) and PEG8000 (blue, lower panel) and after vigorous washing with 50 mM Tris (pH 8) buffer (purple). (G) Injection of high molecular weight PEG crowders leads to reversible vesicle compaction.

A direct quantitative comparison between the effects of molecular crowding seen here on highly curved LUVs and previous work on GUVs is not entirely straightforward. For instance, the optical imaging of GUVs encapsulating molecular crowders as performed previously^32^ only reports on macroscopic changes from a cross-sectional slice across the vesicles. In contrast, the smFRET approach described here provides access to the mean dye separation distance across the entire three-dimensional volume of the vesicle. Nevertheless, both sets of vesicles undergo substantial morphological changes in response to molecular crowders at similar concentrations, and thus our smFRET work on sub-micron sized vesicles is complementary to the optical imaging of GUVs, with both datasets suggesting membrane conformation is strongly influenced by molecular crowding. While the smFRET approach provides details of vesicle perturbation on the nanoscale, a limitation is that it does not quantitatively report on pore formation or solution exchange across the bilayer. Recent studies on GUVs have provided evidence to support the transient formation of nanoscale pores induced by osmotic pressure and variations in membrane tension^54-56^. While we obtained limited evidence for the formation of pores induced by sorbitol (**Figure 3S**), whether this is also true for the sub-micron sized vesicles in response to PEG-based crowders remains an open question. We expect the smFRET approach to play a key role in identifying conformational changes within single vesicles in response to a wide variety of complex crowding conditions, including various solvents, solutes of varying molecular weight, composition, concentration, temperature, pH and viscosity.

## Conclusions

In this work, we characterized the response of synthetic sub-micron sized vesicles to environmental crowding conditions and found that both sorbitol, and polymers of PEG and Ficoll influence their lipid packing density and size, even at modest concentrations of crowder. We conclude that molecular crowding leads to a global compaction of the intact vesicle structure, with our hypothesis supported by the direct observation of single vesicles by electron microscopy. Given the variability of molecular crowding *in vivo*, we subjected the vesicles to a range of molecular weight crowders and unlike the effects of crowding on nucleic acids, which indicate smaller molecules induce more pronounced crowding effects^57,58^, our observations point towards an intriguing size dependence. On the one hand, we observed that crowding by the low-molecular weight sorbitol leads to permanent vesicle compaction via a biophysical mechanism that likely involves integration of the crowder into the vesicle bilayer and membrane dehydration. In contrast, crowding by longer polymers of PEG and Ficoll induced reversible vesicle compaction, likely via an excluded volume effect ^23,59-61^, with the observed structural changes found to be more pronounced with higher molecular weight crowding agents. It should, however, be noted that these observations were made exclusively on phosphocholine-based vesicles, and future work is required to disentangle the influence of lipid composition and cholesterol content. Whether the presented FRET sensor can also be adapted to sense morphological changes to extracellular vesicles *in vivo* has yet to be explored, but we note that morphological compaction has been observed here in synthetic vesicles with similar compositions, sizes and structures.

Understanding the structural stability of sub-micron sized vesicles, especially those with high curvature, and how they dynamically respond to molecular crowders has a number of implications. First, important trafficking pathways between the endoplasmic reticulum rely on the formation of highly curved vesicles, and regulation of extramembrane crowding may be one mechanism through which vesicles alter morphology for vital processes including endocytosis and vesicle budding^62-64^. More generally, crowding-induced changes to local membrane curvature could also play a role in the shaping of organelles, and the triggering of adaptive cellular responses such as gene expression^65^. Second, our measurements indicate that extramembrane crowding leads to variations in the lipid packing density which does not always guarantee stability^66^. In the context of vesicles as drug delivery vehicles, it is therefore possible that molecular crowding influences the efficiency at which encapsulated drug molecules are released and therefore the fate of the targeted cell. Finally, the ability to precisely control vesicle morphology is highly desirable for a number of biotechnological applications^67^. For instance, we foresee that the ability to alter vesicle curvature *in vitro* could open a platform for investigating crowding effects on membrane protein signalling, and for evaluating encapsulation efficiencies in more physiologically relevant conditions.

## Acknowledgements

S. D. Q. acknowledges support from Alzheimer’s Research UK (RF2019-A-001). This study was also supported by BBSRC (BB/W000555/1) and the Leverhulme Trust (RPG-2019-156). We thank Prof. Daniella Barilla (Department of Biology, University of York, UK) for use of DLS instrumentation, Prof. Thomas Krauss (Department of Physics, University of York, UK) for use of SEM Nanocentre facilities, the Bioscience Technology Facility (Department of Biology, University of York, UK) for use of fluorescence spectroscopy apparatus, Dr. Jamie Blaza (University of York, UK) for Cryo-TEM support, and Prof. Marco Fritzsche (University of Oxford, UK) for the generous donation of FliptR.

## Data accessibility

Data associated with this study is freely available from DOI:10.5281/zenodo.7081992.

