## supplementary methods for "Crowding induced morphological changes in synthetic lipid vesicles determined using smFRET"

### SUPPORTING INFORMATION

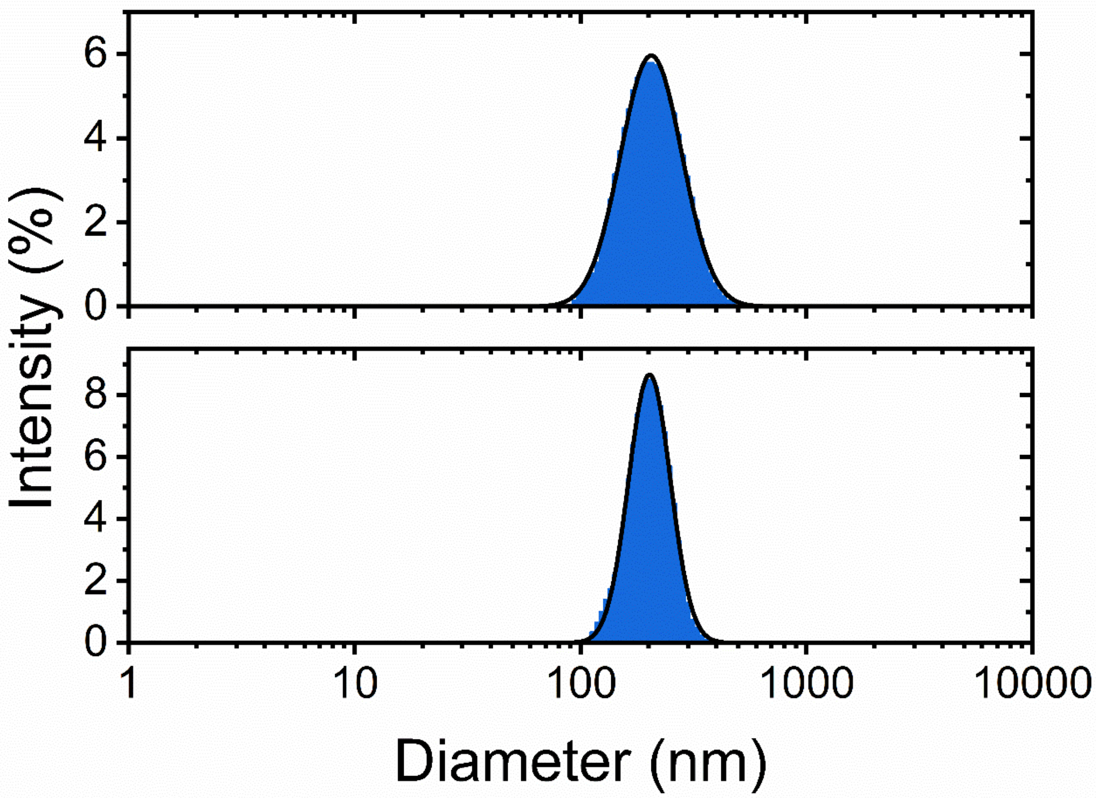


**Figure S1.** Representative DLS diameter distributions obtained from DiI and DiD loaded POPC vesicles (top panel) and unlabelled vesicles (bottom panel). Solution conditions: 50 mM Tris, pH 8.

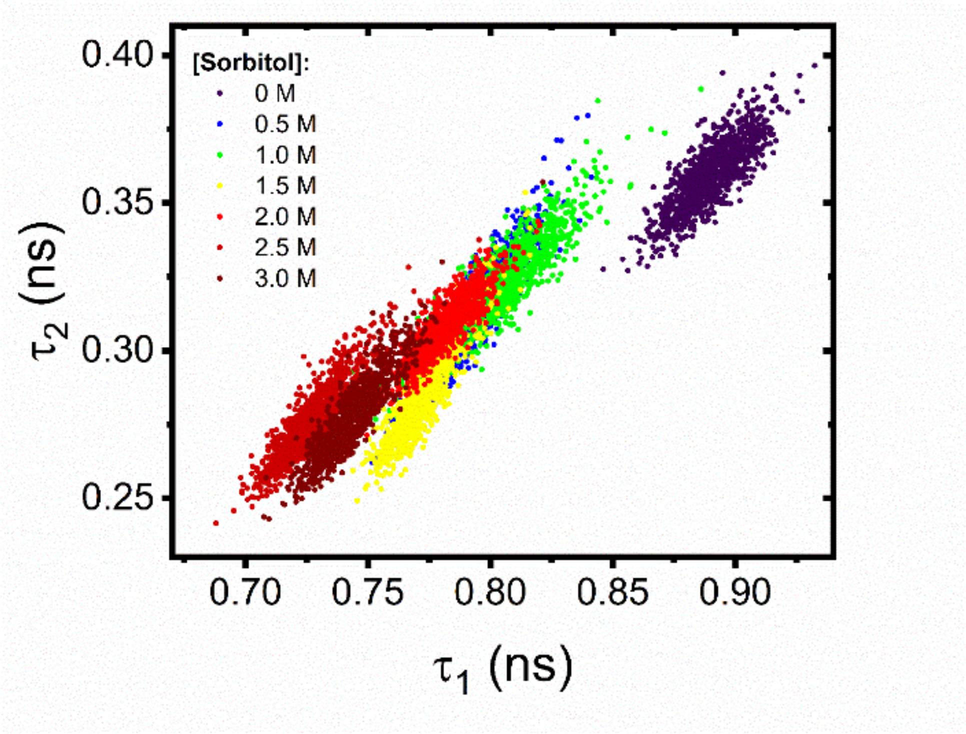


**Figure S2.** Representative variations in the lifetime components $\tau_1$ and $\tau_2$ extracted from DiI and DiD loaded POPC vesicles in the absence and presence of sorbitol.

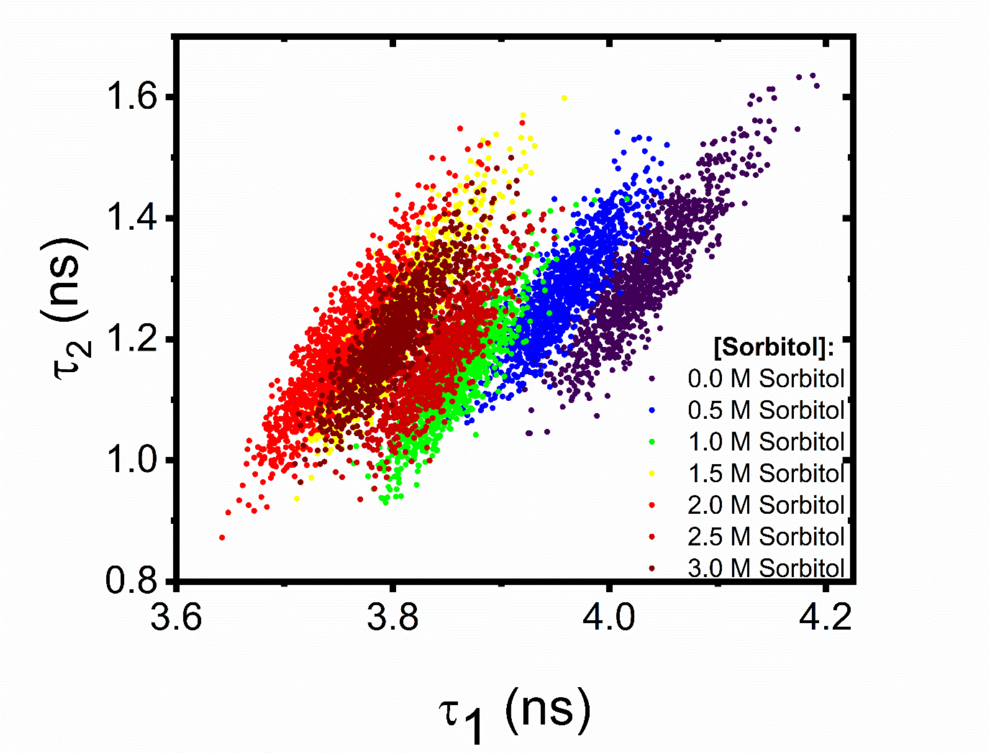


**Figure S3.** Representative variations in the lifetime components $\tau_1$ and $\tau_2$ extracted from FliptR loaded POPC vesicles in the absence and presence of sorbitol.

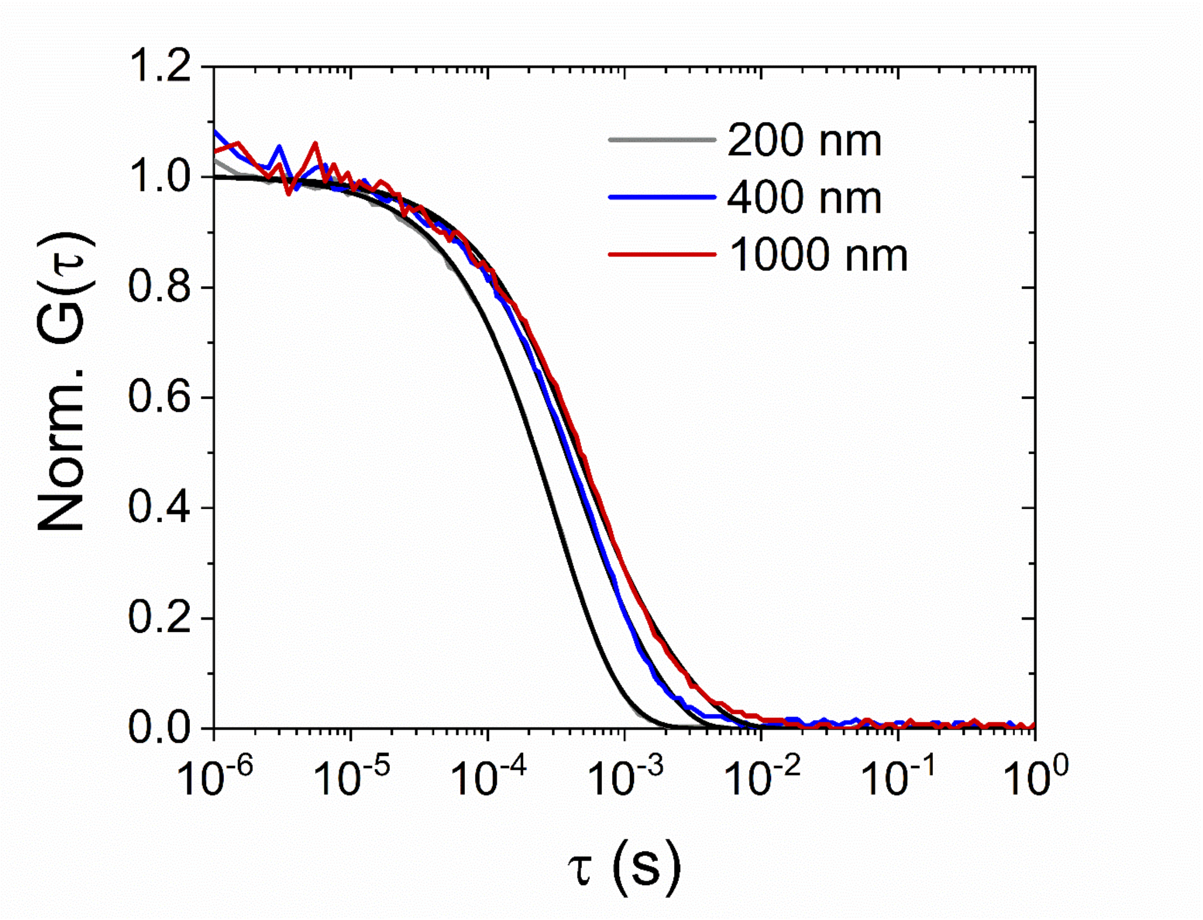


**Figure S4.** Representative variations in the normalized correlation curves obtained by DLS from DiI and DiD loaded POPC vesicles extruded through 200 nm (gray), 400 nm (blue) and 1000 nm (red) pore diameter polycarbonate membrane filters. Solid black lines represent fits as describe in the Methods Section. Solution conditions: 50 mM Tris, pH 8.

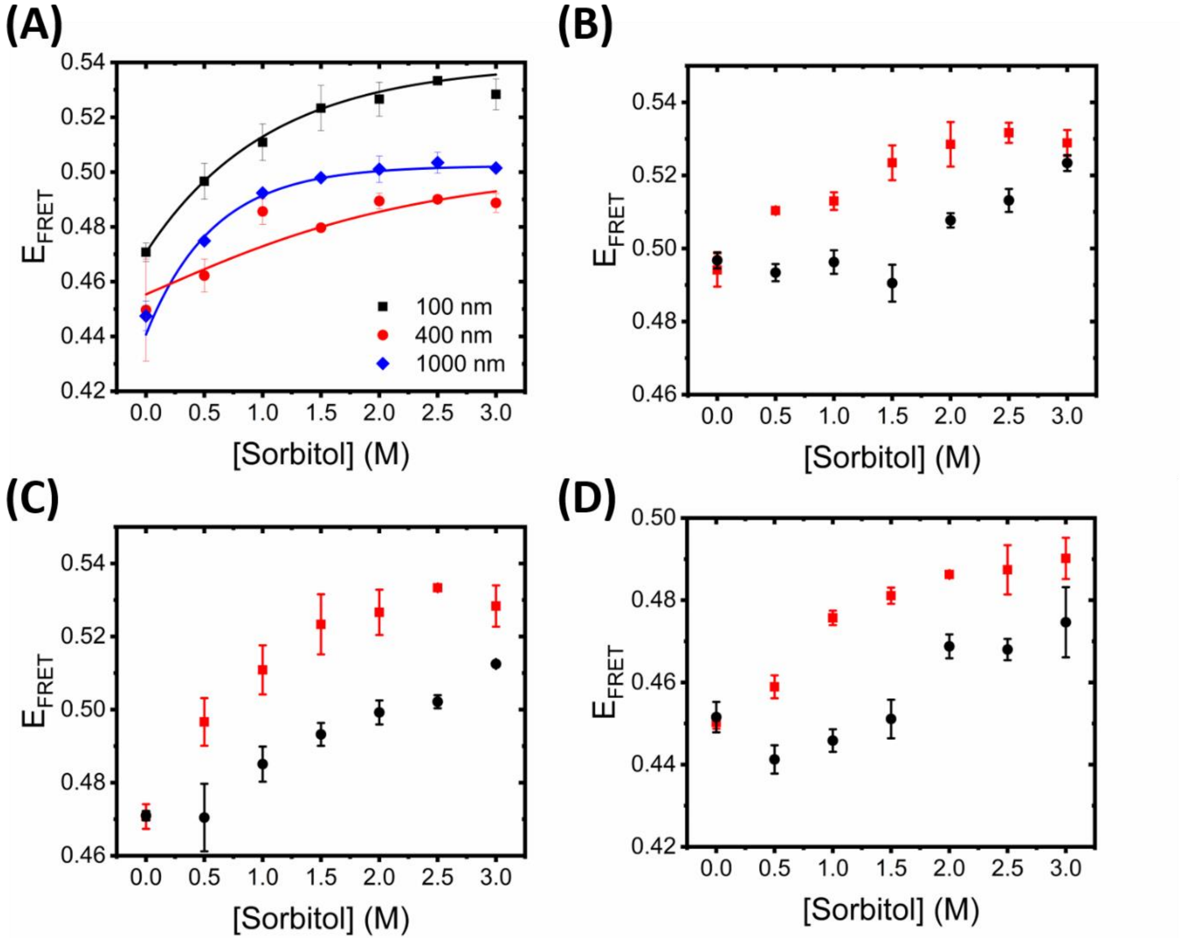


**Figure S5.** (A) Representative variation in $E_{FRET}$ as a function of vesicle size. Also shown are $E_{FRET}$ variations obtained from 100 nm diameter POPC (red) and POPS (black) vesicles at (B) 4$^{o}$C, (C) 21$^{o}$C and (D) 37$^{o}$C.

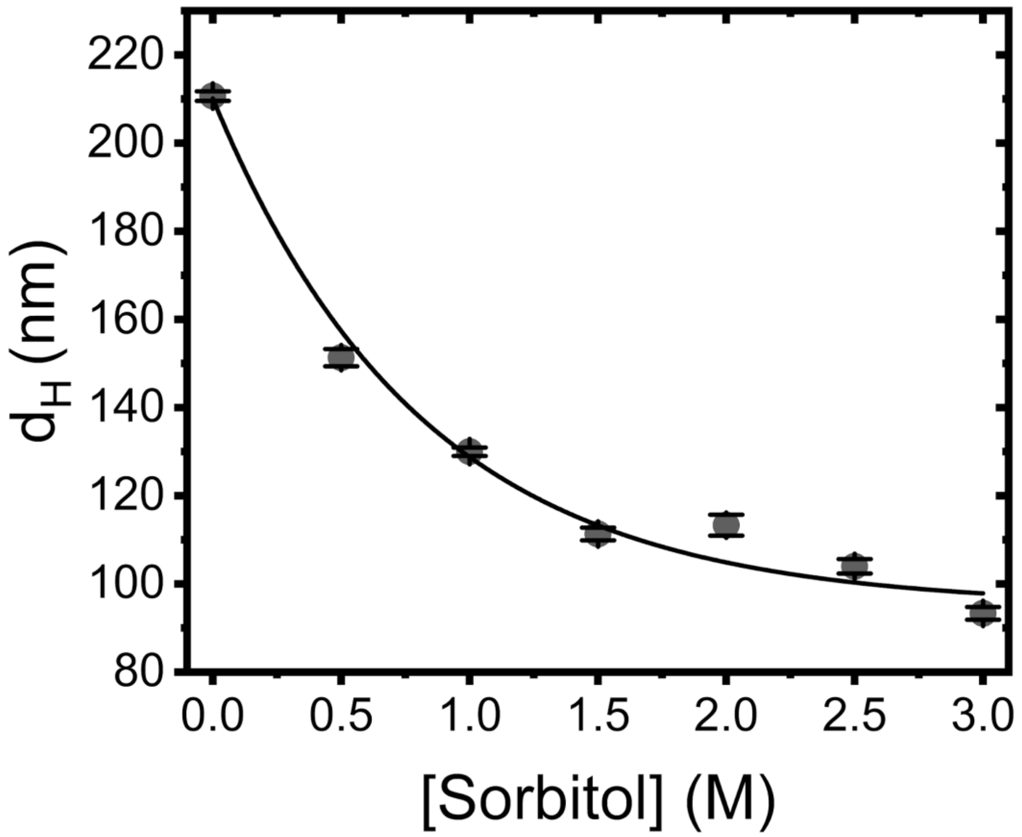


**Figure S6.** Variation in the mean hydrodynamic diameter of POPC-loaded LUVs under crowding conditions.

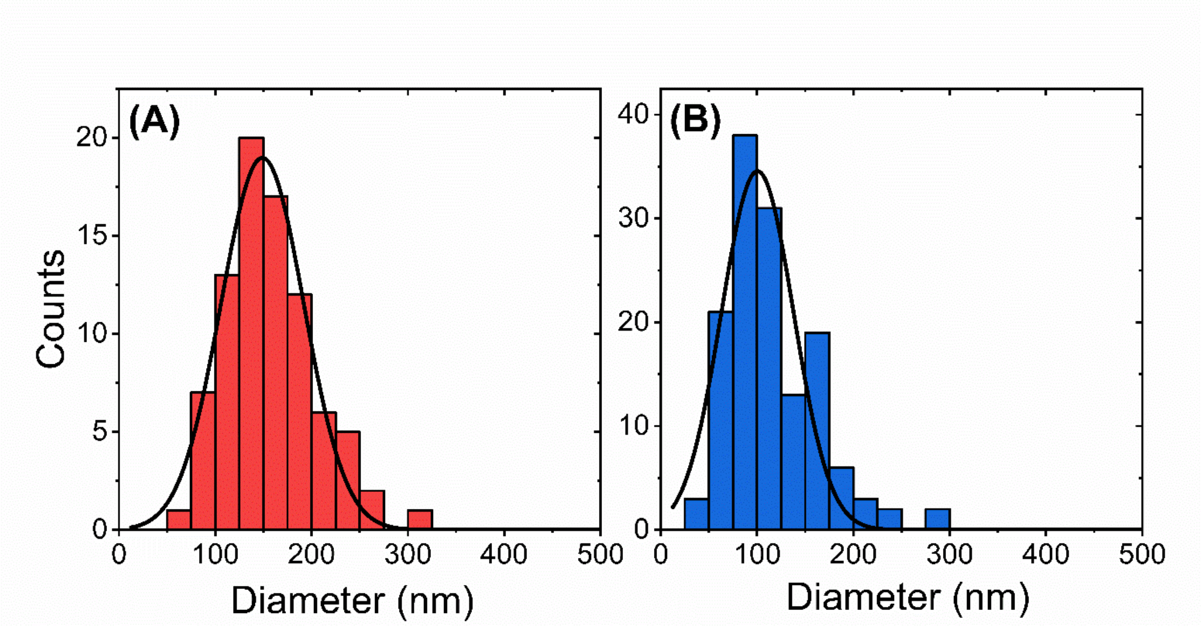


**Figure S7.** Comparison of vesicle diameter distributions obtained by cryo-TEM in (A) the absence (*N* = 84) and (b) the presence of 0.5 M sorbitol (*N* = 138). Solid black lines correspond to Gaussian fits.
